# Enabling functionality and translation fidelity characterization of mRNA-based vaccines with a platform-based, antibody-free mass spectrometry detection approach

**DOI:** 10.1101/2024.05.14.594137

**Authors:** Alyssa Q. Stiving, Benjamin W. Roose, Christopher Tubbs, Mark Haverick, Ashley Gruber, Richard R. Rustandi, Jesse Kuiper, Matthew Schombs, Hillary Schuessler, Xuanwen Li

## Abstract

The success of mRNA-based therapeutics and vaccines can be attributed to their rapid development, adaptability to new disease variants, and scalable production. Modified ribonucleotides are often used in mRNA-based vaccines or therapeutics to enhance stability and reduce immunogenicity. However, substituting uridine with N^1^-methylpseudouridine has recently been shown to result in +1 ribosomal frameshifting that induces cellular immunity to the translated off-target protein. To accelerate vaccine development, it is critical to have analytical methods that can be rapidly brought online to assess the functionality and translation fidelity of mRNA constructs. Here, a platform-based, antibody-free method was developed using cell-free translation (CFT) and liquid chromatography-tandem mass spectrometry (MS) that can detect, characterize, and provide relative quantification of antigen proteins translated from mRNA vaccine drug substance. This workflow enabled the evaluation of mRNA subjected to thermal stress as well as bivalent (i.e., two mRNA encoding different antigen variants) drug substance. Additionally, the MS detection approach exhibited high sensitivity and specificity by accurately identifying all six translated proteins and their relative abundances in a dose-dependent manner following transfection of human cells with a hexavalent mRNA mixture encapsulated in lipid nanoparticles (LNPs), despite significant protein sequence homology. Expanding on these efforts, we show the utility of the CFT-MS approach in identifying the presence and junction of +1 ribosomal frameshifting resulting from N^1^-methylpseudouridation. Overall, this CFT-MS methodology offers a valuable analytical tool for the development and production of mRNA-based vaccines by facilitating the evaluation of mRNA quality and functionality while ensuring accurate translation of antigen proteins.

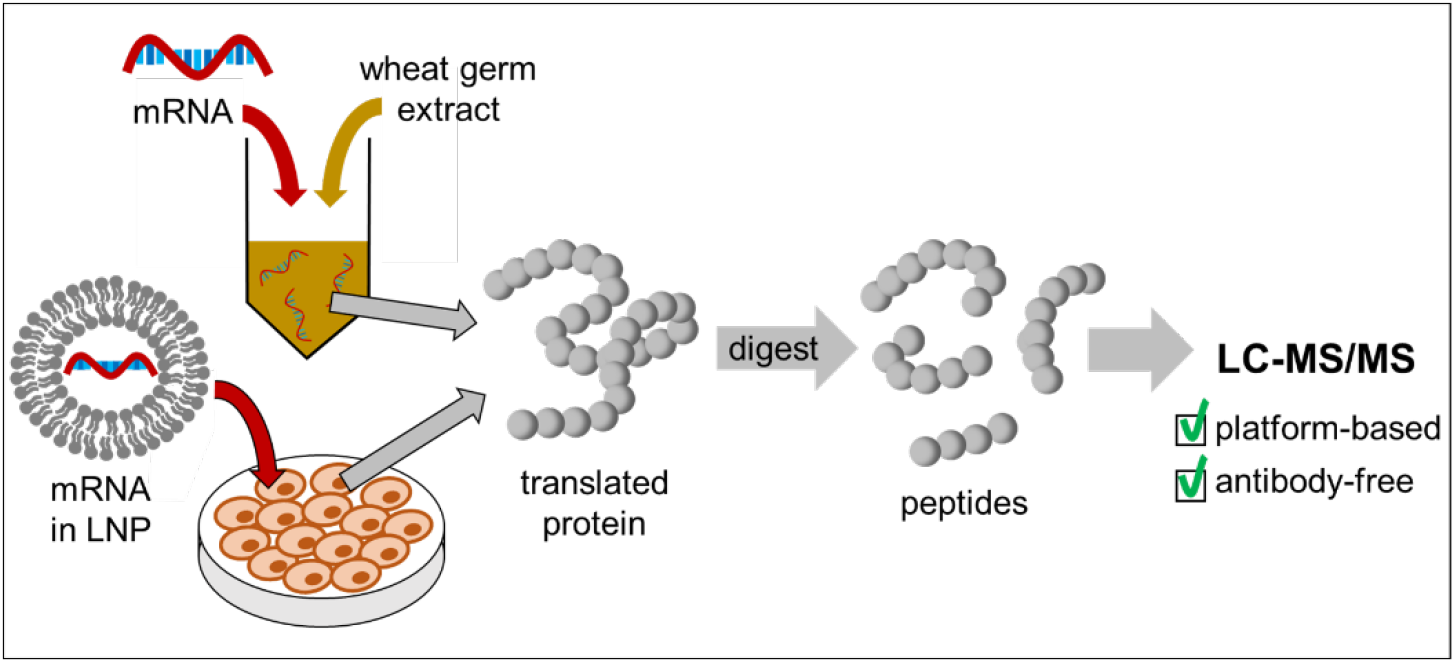

## Introduction

The success of the Pfizer-BioNTech and Moderna mRNA vaccines against severe acute respiratory syndrome coronavirus disease 2019 (SARS-CoV-2) has placed the spotlight on mRNA as an appealing platform for vaccinating against a wide range of diseases (e.g., influenza^1^, HIV^2^, and RSV^3^).^4,5^ mRNA vaccines often consist of lipid nanoparticles (LNP) encapsulating mRNA encoding the antigen sequence which, when translated by host cells, leads to antigen presentation and stimulation of immune response. Key advantages of mRNA vaccines include ease of manufacture, amenability to updating the antigen sequence to match rapidly mutating pathogens (e.g., SARS-CoV-2), and the ability to multiplex different mRNA constructs within the same vaccine product.^6^ The use of modified ribonucleotides is common in mRNA-based therapeutics or vaccines because they decrease innate immunogenicity and increase mRNA stability, leading to the recognition of these advancements with the 2023 Nobel Prize in Physiology or Medicine.^7–10^ As an example, the clinically approved SARS-CoV-2 mRNA vaccines incorporate N^1^-methylpseudouridine to achieve these attributes.^11,12^ Using *in vitro*-transcribed mRNA as a therapeutic or vaccine is reliant on accurate translation of the encoded antigen protein following transfection into target cells, yet the use of modified ribonucleotides and the corresponding impact on mRNA translation fidelity were, until recently, not thoroughly investigated.

Recent work by Mulroney and coworkers investigated the impact that N^1^-methylpseudouridylation has on mRNA translation integrity and demonstrated that this modification results in +1 ribosomal frameshifting in vitro, albeit at significantly lower abundance than the in-frame translated protein.^13^ This was observed along with cellular immunity in both mice and humans to the +1 frameshifted products from the BNT162b2 (SARS-CoV-2) vaccine (trade name: Comirnaty; generic name: tozinameran) mRNA following vaccination.^13^ This +1 ribosomal frameshifting occurred in one of the regions containing a “slippery sequence” and the aforementioned work by Mulroney, et al. suggests that optimizing slippery sequences is a proactive and effective strategy for reducing the translation of frameshifted products. Despite no adverse outcomes being reported due to the incorrect translation of these vaccines associated with frameshifted products, this serves as a cautionary tale to optimize the mRNA sequence to prevent off-target effects for future mRNA therapeutics and vaccines.

To support the development, optimization, and production of mRNA vaccines, a wide range of analytical methods have been developed to better understand and monitor critical quality attributes (CQAs) of both the mRNA itself, the LNP, and the mRNA-LNP complex.^14,15^ For mRNA, examples of these CQAs include sequence identity, 5’-capping efficiency, 3’ poly(A) tail length, and mRNA integrity (i.e., intactness) – all properties that ultimately affect the translation of antigen protein and, therefore, vaccine efficacy. To complement these methods for characterizing specific attributes of the mRNA, it is also important to have methods for evaluating the overall functionality of the mRNA, defined here as the successful translation of antigen by the ribosome.^16^ To this end, we developed an analytical assay to assess mRNA functionality based on cell-free translation (CFT), also known as *in vitro* translation (IVT) or cell-free protein synthesis (CFPS) in the literature.^17,18^ Unlike cell-based expression systems that require cell culture over multiple days, CFT is capable of rapidly assessing mRNA functionality as protein translation is observed within just a couple of hours. Moreover, CFT is performed by delivering mRNA directly to the ribosomal machinery, whereas in cell-based expression systems, measurement of mRNA functionality can be confounded by the quality of the LNP used to deliver the mRNA into the cells. Several CFT systems are commercially available, derived from bacteria, yeast, plant, insect, and mammalian cells.^17–19^ Of these, wheat germ extract (WGE) is well established in the literature and is reported to have the highest yields of any eukaryotic CFT system^20–22^, thus making it the logical starting point for the development of a CFT assay to assess mRNA functionality. While CFT has a role in providing a rapid means for assessing mRNA functionality without the confounding effects of the LNP delivery system, cell-based translation (CBT) may be used for the assessment of mRNA drug product formulated in LNPs and often provides a more representative output of mRNA functionality *in vivo*. For these reasons, we also assessed the ability to characterize mRNA quality and functionality using CBT.

To quantitate the amount of antigen translated by WGE-CFT, an automated capillary Western blot method, Simple Western™ (SW), was chosen for initial method development. SW consists of protein separation by capillary electrophoresis (CE) followed by immobilization of the separated proteins to the capillary wall and immunodetection with a suitable primary antibody. SW measures both the molecular weight and relative signal of immunodetected proteins. However, it does not provide sequence-level information, thus limiting its specificity and ability to detect frame-shifting events during antigen translation from mRNA. A significant drawback of SW is its reliance on antigen-specific primary antibodies as a critical reagent. For multivalent vaccines, which may contain mRNA encoding variants of the same antigen with high levels of similarity, it may prove challenging to develop antibodies that are highly specific to each antigen. Additionally, if the mRNA antigen sequence is updated in response to newly emerged or seasonal serotypes, antibody redevelopment may be required which can be both time consuming and costly.

To overcome the potential limitations of using an immunodetection method, we sought to demonstrate the viability of using liquid chromatography-tandem mass spectrometry (LC-MS/MS, or “MS” as used throughout this manuscript) as an antibody-free alternative for the identification and relative quantification of antigen produced by CFT or CBT of mRNA. Bottom-up MS involves subjecting a mixture of proteins to enzymatic digestion and analyzing the resulting peptide mixture by MS. The resulting spectra can then be matched against a protein database that has been “digested” *in silico* using the enzyme of choice for identification of proteins in the sample. Bottom-up MS is a highly specific and sensitive technique, allowing for distinction between proteins even with high sequence homology without relying on specific antibodies. This enables a antibody-free platform-based approach to protein detection and characterization.

Diseases that mutate rapidly necessitate expedited vaccine development to provide relevant and often seasonal protection. Examples include COVID-19 (SARS-CoV-2) and influenza. Responding to such diseases requires analytical assays that are readily available and don’t require months of development lead time. Because of the rapid mutation of SARS-CoV-2, Delta (B.1.617.2) spike protein and Omicron spike protein mRNA constructs serve as ideal examples to demonstrate the utility of such methods. Here, we demonstrate the utility of this platform-based MS method in detecting, characterizing, and providing relative quantification of proteins translated from CFT of mRNA. We also evaluated platform-based MS detection in a proof-of-concept experiment wherein MS was used to detect and provide relative quantification of proteins translated from a hexavalent mRNA drug product transfected in human cells using CBT. Furthermore, we demonstrate that using a CFT-MS approach, we can interrogate +1 ribosomal frameshifting occurring from N^1^-methylpsuedouridylation of the mRNA and enable identification of the frameshifting junction. The resulting workflow is a sensitive, antibody-free, platform-based approach that can be used to confirm identity and provide relative quantification of protein(s) translated from mRNA from either CFT or CBT for the assessment of mRNA quality, functionality, and translation fidelity.

## Methods

### Preparation of mRNA

Prior to handling mRNA, all bench surfaces, pipettes, and gloves were wiped with RnaseZap™ (Invitrogen, cat. no. AM9786) followed by 70% isopropyl alcohol. mRNA was handled using only RNase-free consumables. mRNA was diluted in 1 mM sodium citrate pH 6.4 (Thermo Fisher Scientific). The concentration of mRNA was measured using a NanoDrop spectrophotometer (Thermo Fisher Scientific) blanked with mRNA diluent.

### Cell-free translation (CFT) of mRNA

Cell-free translation (CFT) was performed using wheat germ extract (Promega) supplemented with Complete Amino Acid Mixture (Promega), potassium acetate (Promega), RNasin Plus RNase inhibitor (Promega), and nuclease-free water (Thermo Fisher Scientific). mRNA was added to the CFT mixture and incubated in a heat block at 25 °C for 2 h. After 2 h, the CFT mixture was aliquoted for subsequent analysis by Simple Western™ or mass spectrometry. Aliquots for Simple Western™ were diluted 10-fold in 1X Simple Western™ Sample Buffer prior to freezing at −70 °C, whereas aliquots for LC-MS/MS were stored neat at −70 °C.

### Cell-based translation (CBT) of mRNA

Cell-based translation (CBT) was performed using Huh7 cells prepared in a 96-well plate at 85-95% confluency. Cells were transfected with a hexavalent mRNA drug product (DP) comprised of mRNAs 1 through 6 that encode for various proprietary spike protein sequences encapsulated in proprietary lipid nanoparticles (LNPs) for delivery. Each mRNA DP was transfected at a dose of 6.67 or 4.44 ng per mRNA construct and incubated at 37°C for 18 hours. After incubation, the supernatant was removed and the cell monolayer was gently washed three times with approximately 200 µL PBS. Following this, the cells were lysed with the addition of 100 µL of cell lysis buffer from the EasyPep™ MS Sample Prep kit (Thermo Fisher Scientific, Waltham, MA). After mixing vigorously (10-15X), the collected cell lysate containing the translated proteins was frozen prior to preparation for mass spectrometry analysis as outlined below.

### Automated Western blotting

Automated capillary Western blotting (i.e., Simple Western™) of CFT samples were performed using either a Wes or Jess instrument (ProteinSimple) following the manufacturer’s procedure. Briefly, CFT samples were denatured and reduced by treatment with Master Mix solution prepared from the EZ Std kit Pack (ProteinSimple) followed by heating at 95 °C for 5 min. Simple Western was then performed using Antibody Diluent 2 (ProteinSimple) as the blocking reagent, anti-CoV spike S2 mouse monoclonal antibody (Sino Biological) as the primary antibody, and anti-mouse IgG HRP conjugate (ProteinSimple) as the secondary antibody. Immunoprobe detection was performed using a 1:1 mixture of Luminol-S (ProteinSimple) and Peroxide (ProteinSimple) and measuring chemiluminescence. All data analysis was performed in Compass software (ProteinSimple).

### Microchip capillary electrophoresis

The integrity of thermally-degraded Delta mRNA was measured using microchip capillary electrophoresis (MCE) as described previously.^23^ Briefly, mRNA samples were prepared in singlet by diluting to a final concentration of 10 µg/mL in 10% (w/v) Brij® 58 (Acros Organics) and 1X Sample Buffer (Perkin Elmer) in formamide (Sigma-Aldrich). The treated samples were then heated at 70 °C for 10 minutes followed by incubation on ice for at least 5 minutes. The samples were analyzed in duplicate by the RNA Standard Sensitivity Assay on a LabChip GXII Touch (Perkin Elmer) using a DNA 5K/RNA/CZE LabChip. The mRNA integrity of each sample was calculated by dividing the peak area of intact mRNA by the total peak area of the sample.

### Mass spectrometry

Cell-free translation (CFT) or cell-based translation (CBT) mRNA samples were prepared as described above and the resulting translated protein(s) were thawed then sonicated for 5 minutes. Following this, the contents of each sample were digested using a suspension trap (S-trap™ micro, Protifi, Fairport, NY) following manufacturer recommendations. Briefly, samples were combined with sodium dodecyl sulfate (SDS, EMD Millipore) to a final concentration of 5% SDS, proteins were reduced with dithiothreitol (Thermo Pierce) and then alkylated with iodoacetamide (Thermo Pierce). Next, samples were acidified with the addition of phosphoric acid then combined with S-trap binding buffer (100 mM triethylammonium bicarbonate buffer, 90% methanol, pH 7.1) and loaded onto the S-trap™ columns with centrifugation. After proteins were trapped on the S-trap column, trypsin was added (1 µg per column) and columns incubated at 37°C overnight. Following digestion, peptides were eluted from the S-trap™ and subsequently dried using a SpeedVac followed by reconstitution in 0.1% formic acid in water. Peptides were subsequently cleaned up with C18 tips (Thermo Pierce), concentrated to dryness, and reconstituted with 0.1% formic acid in water. LC-MS/MS analysis was performed on a nanoAcquity LC (Waters) and Orbitrap Exploris™ 480 mass spectrometer (Thermo Fisher Scientific) with an Easy-Spray HPLC column (ES900, Thermo Fisher Scientific) and 170 min gradient. Alternatively, peptides eluted from the S-trap™ were cleaned up directly by loading onto EvoTips and analyzed by LC-MS/MS using an Evosep One LC and Thermo Exploris 480. It is worth noting that the sonication and C18 cleanup steps were found to be critical for avoiding column clogging due to the viscous nature of the wheat germ extract. Samples analyzed with the Evosep One used an EV1106 column and the 15SPD (15 “samples per day”, corresponding to an 88 min) predefined gradient. Mobile phases consisted of 0.1% formic acid in water (A) and 0.1% formic acid in acetonitrile (B) in both LC setups. All MS/MS data was collected in data-dependent acquisition for all experiments unless noted otherwise. The spray voltage was set to 1.9 kV, with the ion transfer tube was to 275°C. The full scan range was 275-1200 *m/z*, MS1 resolution = 60,000, MS2 = 15,000. The normalized AGC target for MS1 = 250% with a maximum injection time of 20 ms. MS2=50% with a maximum injection time set to auto. Number of dependent scans was 30. Following untargeted identification of the +1 frameshifted Delta (B.1.617.2) spike protein peptide GYHLMSFPR, a targeted MS analysis was used to confirm the observation where the 2+ and 3+ charge states were added to a targeted inclusion list (no retention time window was specified). For this targeted experiment, MS2 AGC 1e5 and maximum injection time 120 ms were used. Resulting RAW data from untargeted experiments were analyzed using Proteome Discoverer 2.5 (Thermo Scientific) with Sequest HT search against the respective proteome (Triticum aestivum, UniProt proteome ID: UP000019116) and expected mRNA translated protein sequences. Proteome Discoverer was searched with a precursor mass tolerance of 10 ppm, fragment mass tolerance of 0.02 Da, up to 1 missed cleavage, and 1% FDR target; results were reported with protein abundance defined as MS1 areas of all identified peptides, which was used to calculate relative abundances throughout this manuscript. For peptide mapping, Genedata Expressionist 17.0 was used with a standard peptide mapping workflow including the expected mRNA translated protein sequence and allowing for up to one missed cleavage.

## Results & Discussion

As a means of generating translated protein to assess mRNA functionality (i.e., translation of antigen protein), we utilized two approaches: cell-free translation (CFT) using wheat germ extract and cell-based translation (CBT) using Huh7 cells. Each mRNA translation approach has its own advantages as discussed previously. Detection of translated protein(s) following CFT is often accomplished using methods such as radiolabeling, immunoprobe (e.g., Western blotting, enzyme-linked immunosorbent assay (ELISA)), or functional assays like Luciferase activity readout.^24^ Among these methods, we selected automated Western blotting by Simple Western™ (SW) due to its relatively high throughput, automation, reproducibility, and availability of antibodies for antigen detection.^25–27^ However, the CFT-SW approach still relies on suitable antibodies for antigen detection, which can pose challenges including increased assay development time and cost for antibody screening, high detection limits and variability when using poor quality antibodies, the need for alternative detection assays if suitable antibodies cannot be found, and limited ability to multiplex detection for analysis of mRNA mixtures due to antibody cross-reactivity. Mass spectrometry, on the other hand, is a highly sensitive technique that lends itself well to a platform-based approach for protein detection and relative quantification, alleviating the need to perform antibody screening and re-development for each mRNA construct or antigen target. For this reason, we coupled MS to both CFT and CBT approaches. These workflows are outlined in Figure 1.

**Figure 1.**
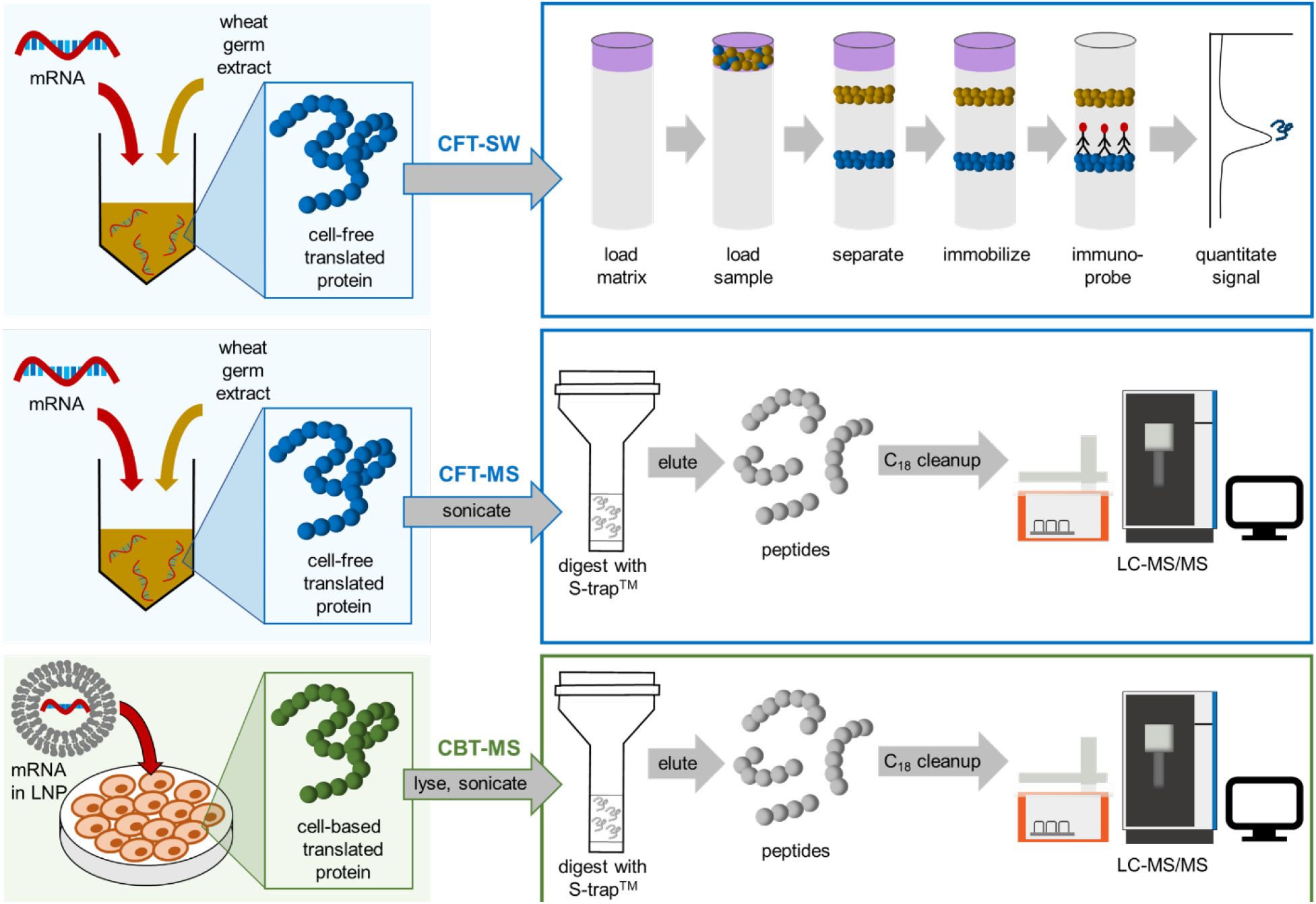
Scheme outlining the cell-free and cell-based translation workflows coupled to Simple Western™ (CFT-SW) or reversed-phase liquid chromatography tandem mass spectrometry (CFT-MS and CBT-MS) workflows for detection of proteins translated from mRNA.

### Assessment of mRNA functionality by detecting translated protein identity and sequence confirmation by CFT-MS

Another key advantage of mass spectrometry-based detection of proteins is the ability to characterize the primary structure of the translated protein(s). This enables confirmation of mRNA identity and quality as assessment of the translated protein sequence can ensure that the mRNA encodes for the correct protein sequence and has an appropriately located stop codon. Following the initial development of the CFT-MS workflow, assay performance was evaluated to demonstrate this capability. To accomplish this, mRNA encoding for the SARS-CoV-2 Delta (B.1.617.2) spike protein was analyzed using the CFT-MS workflow where protein digestion was accomplished with trypsin, chymotrypsin, or alpha lytic protease. While trypsin can be selected as a broadly applicable enzyme for a platform-based approach that allows for protein detection, relative abundance, and primary sequence confirmation, the depth of sequence coverage is dependent on the protein sequence of interest. Therefore, the use of additional enzymes suited to the protein sequence can help improve the sequence coverage, if desired. We analyzed the resulting data from CFT-MS analysis using a peptide mapping approach to confirm the sequence of the SARS-CoV-2 Delta (B.1.617.2) spike protein (Figure 2). Results show that when using these three digestion enzymes, we obtained 94% sequence coverage of this spike protein using MS/MS matches. If mass-only matches are also considered, the sequence coverage increases to 99%. This demonstrates the advantage of using CFT-MS to ensure that the mRNA codes for the expected protein sequence. Additionally, coverage of the N-terminus and C-terminus advantageously enables confirmation that the entirety of the mRNA open reading frame is translated, and the stop codon is in the correct location. If high throughput and accurate quantitation of translated protein without complete sequence confirmation is desired, CFT-MS could be performed with targeted MS analysis that tracks and quantifies selected surrogate peptides, such as with multiple reaction monitoring (MRM) based isotope dilution mass spectrometry (IDMS).^28^ The use of wheat germ extract for CFT simplifies MS analysis as it does not result in glycosylation of the translated protein due the lack of necessary cell machinery in the reaction mixture.^22^ The use of alternative CFT systems, such as HeLa lysate, can be used to assess glycosylation patterns on the antigenic protein as the presence of host glycans can greatly impact immune response.^29^

**Figure 2.**
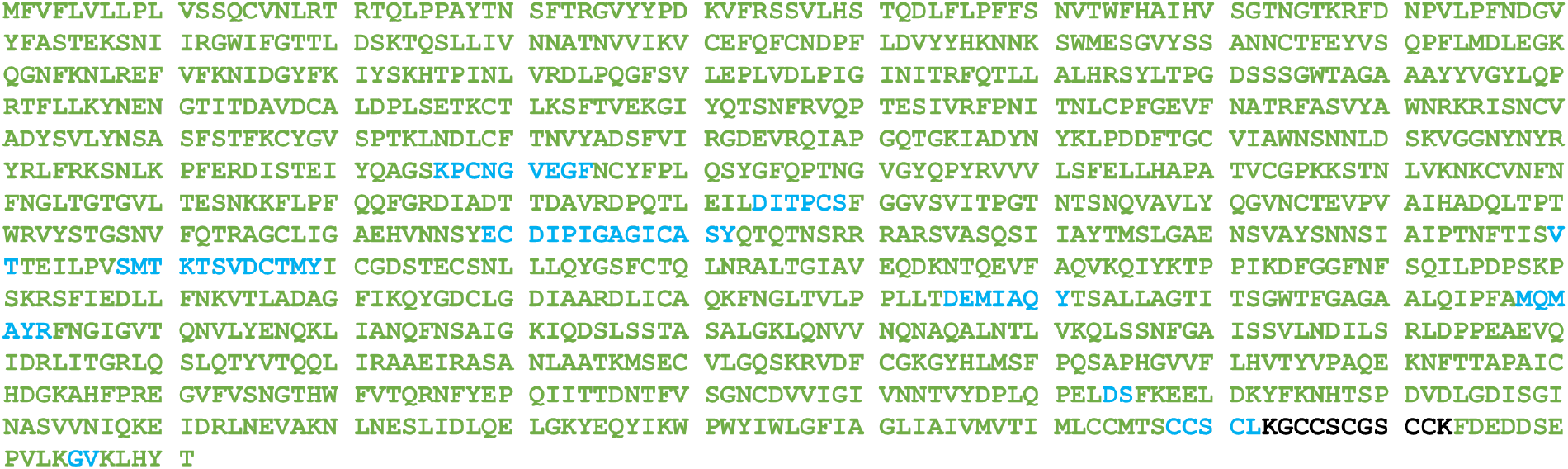
Sequence coverage map from CFT-MS analysis of SARS-CoV-2 Delta (B.1.617.2) spike protein demonstrating the ability of the assay to confirm translated protein identity. Amino acids confirmed with MS/MS matches are represented in green and amino acids identified with mass only matches are represented in blue. Black amino acids indicate no mass only or MS/MS matches were identified. Overall, 94% sequence coverage was obtained from CFT-MS analysis with 13.1 µg/mL mRNA and considering only MS/MS matches. Both the N-terminus and C-terminus were covered with MS/MS matches, allowing verification of correct translation through the entire protein sequence. Peptide mapping results were obtained by combining results from CFT-MS analysis of Delta (B.1.617.2) spike protein using the enzymes trypsin, chymotrypsin, and alpha lytic protease. Additional details regarding which peptides were identified by each enzyme are included in Supplemental Information Figure S1.

### Assessment of mRNA functionality by detecting mRNA dose dependent protein translation by CFT-SW and CFT-MS

Next, we translated SARS-CoV-2 Delta (B.1.617.2) spike protein mRNA *via* CFT at varying concentrations of mRNA in order to evaluate the relationship between mRNA concentration and protein translation by CFT. The relative abundance of translated protein was measured by both SW and MS to compare the two workflows (Figure 3). Results between the two workflows are remarkably similar, showing a strong linear response between 0 and 13.1 µg/mL mRNA, maximum signal at 52.4 µg/mL mRNA, and signal decrease at higher concentrations of mRNA. While initial analysis by CFT-SW was done with a highly sensitive antibody (“antibody 1”), Simple Western™ was repeated using a less sensitive antibody (“antibody 2”) to verify that the maximum observed at 52.4 µg/mL mRNA was not due to detector saturation (results included in Supplementary Information Figure S2 and Figure S3). Because the observed mRNA concentration:antigen translation profile is the same regardless of antibody sensitivity or protein detection technique, these findings suggest that there is a diminishing return to increasing the mRNA concentration when it comes to cell-free translation. The slight decrease in translated protein when mRNA concentration exceeds 52.4 µg/mL is possibly due to mRNA aggregating at higher concentrations, as this loss of protein translation has been observed to varying degrees from other mRNA constructs (data not shown). Additionally, results shown in Figure 3 demonstrate the repeatability of both CFT-SW and CFT-MS workflows. Ultimately, this dose curve demonstrates the ability of both CFT workflows to generate accurate and repeatable relative abundance information about proteins translated from mRNA.

**Figure 3.**
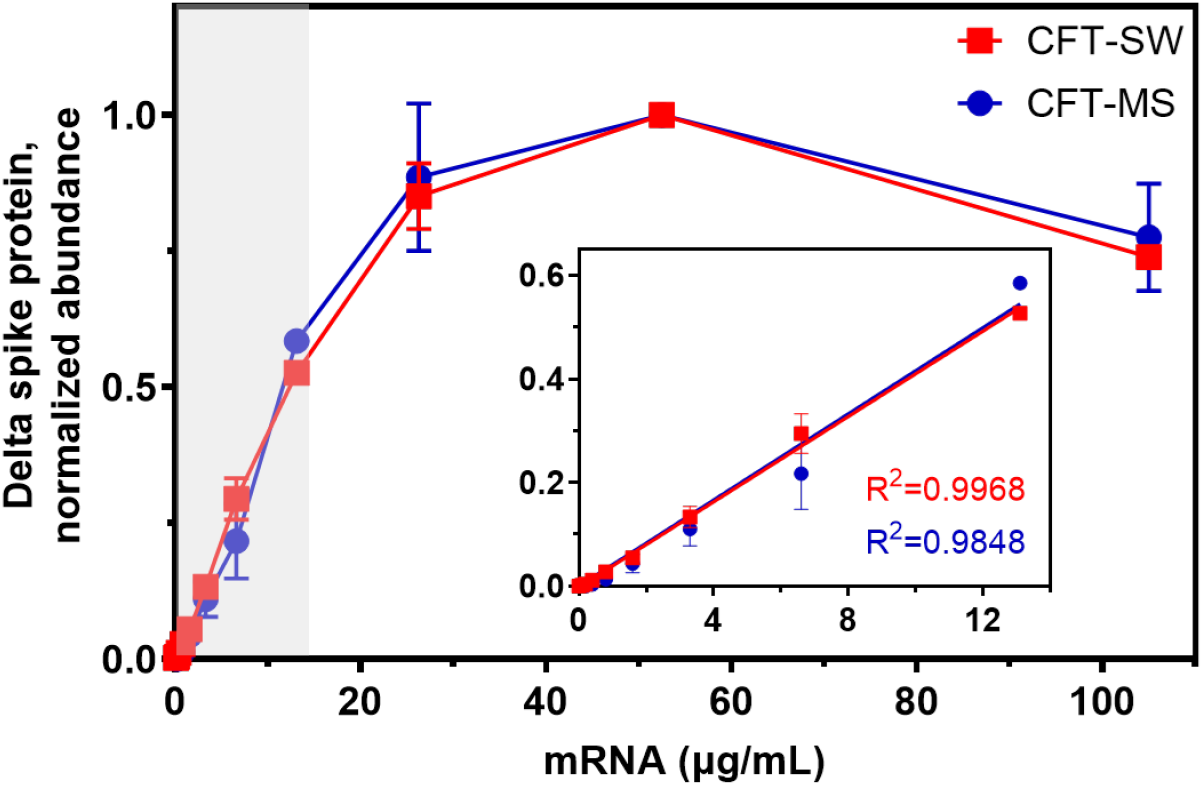
Dose response curve showing the CFT-SW normalized peak area response for CFT-SW (red) and normalized MS1 area response for CFT-MS (blue) detecting translated SARS-CoV-2 Delta (B.1.617.2) spike protein as a function of mRNA concentration. Insets show the linear range for each workflow as highlighted by the gray shaded box, demonstrating strong linearity (R^2^>0.98) in the region of 0 to 13.1 µg/mL mRNA. Plots that show the actual (not normalized) peak area (CFT-SW) and MS1 area (CFT-MS) of translated Delta (B.1.617.2) spike protein are included in the Supplementary Information (Figure S3). Error bars are shown as the standard deviation from duplicate measurements; where error bars are not visible, the error is small enough to be obscured by the data point size.

### Assessment of mRNA quality using CFT-SW and CFT-MS as stability-indicating assays

Process development and stability assessment rely on assays that provide a functional readout of mRNA quality. To interrogate the ability of the CFT-SW and CFT-MS workflows to provide such information, we incubated SARS-CoV-2 Delta (B.1.617.2) mRNA at 50 °C and sampled at various time points up to 55 hours. mRNA integrity at each time point was measured by microchip capillary electrophoresis (MCE) (Figure 4).^23^ Additionally, we assessed the functionality of mRNA at each time point by wheat germ CFT combined with either SW or MS detection (Figure 4). Results from both CFT-SW and CFT-MS demonstrate that the amount of translated protein (relative to the non-degraded time point t=0) decreases accordingly as the mRNA is degraded. These results indicate that the relative abundance of translated protein from CFT-SW and CFT-MS workflows can indeed be used as a stability-indicating assay for the assessment of mRNA quality and functionality. If absolute quantification of the fully translated protein is required, a mass spectrometry approach that targets the C-terminus to ensure complete protein translation could be employed using single reaction monitoring (SRM) or multiple reaction monitoring (MRM).

**Figure 4.**
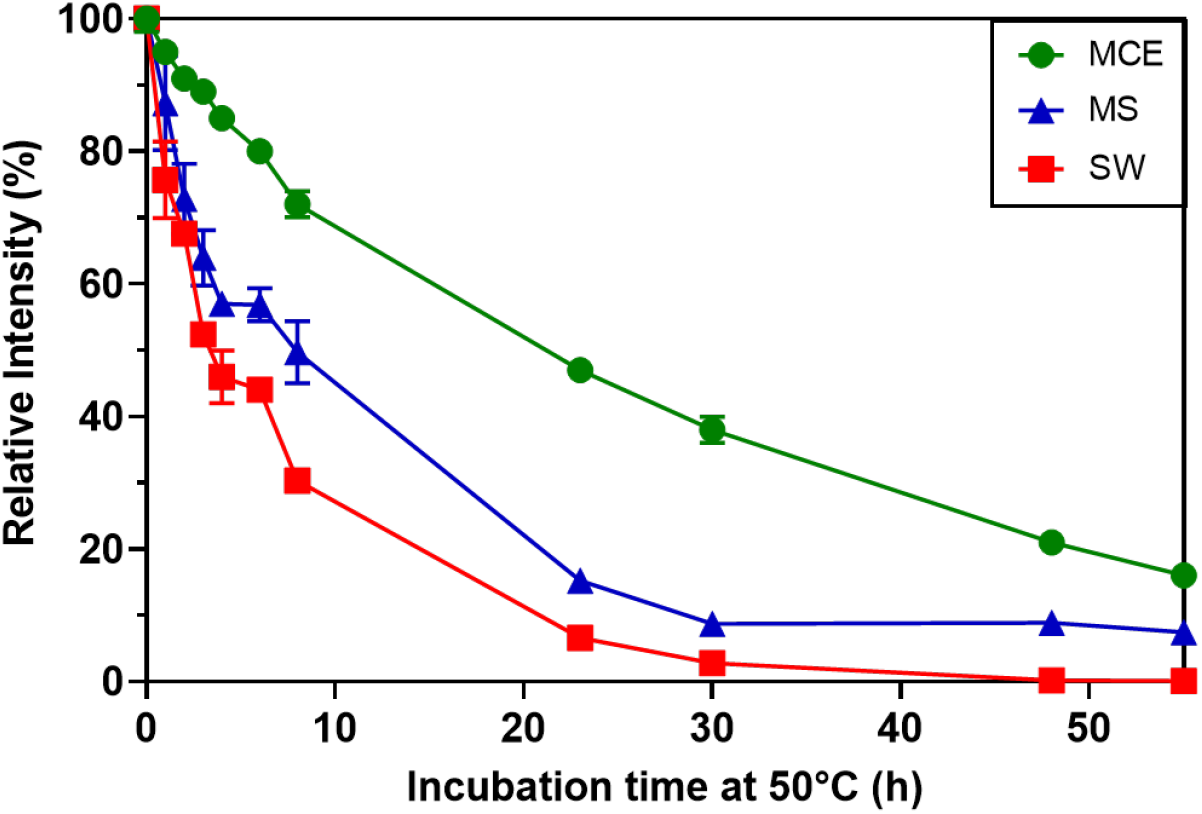
Demonstration of the functional readout from CFT-SW and CFT-MS assays. Following incubation of SARS-CoV-2 Delta (B.1.617.2) mRNA at 50°C for varying time points to degrade the mRNA, wheat germ CFT was performed on 2 µg/mL mRNA from each time point. Translated protein was detected by either Simple Western™ (“SW”, red squares) or LC-MS/MS (“MS”, blue triangles). The mRNA itself was also analyzed by microchip capillary electrophoresis as a direct measurement of mRNA integrity (“MCE”, green circles, electropherograms provided in Figure S4). The y-axis shows relative intensity in percent corresponding to: mRNA integrity normalized to t=0 for MCE, chemiluminescence signal relative to t=0 for SW, and MS1 area relative to t=0 for MS. Error bars are shown as the standard deviation from duplicate measurements; where error bars are not visible, the error is small enough to be obscured by the data point size.

### Assessment of mRNA functionality by evaluating multi-valent mRNA mixtures with high translated protein sequence homology by CFT-SW and CFT-MS

The adaptation of viruses to evade immune response creates variants that may require rapid revision of vaccine development to target the appropriate antigen. To complicate this further, several variants of the same virus may be in circulation at a given time. One such approach to tackle these challenges is to generate a multi-valent vaccine aimed at targeting several variants of the virus of interest. While multi-valent mRNA vaccines have already proven helpful in the fight against COVID-19^30^, an increase in valency can create analytical challenges, especially if the encoded antigens share high sequence similarity. For example, the spike proteins of the Delta (B.1.617.2) and Omicron (B.1.1.529) strains of SARS-CoV-2 share over 97% sequence homology. As such, selective immunodetection of antigens translated from a bivalent Delta+Omicron vaccine drug substance demonstrates a significant analytical challenge.

Even with the example shown here, extensive screening was needed to identify SARS-CoV-2 antibodies specific to Delta (B.1.617.2) and Omicron (B.1.1.529) spike protein (and even the Omicron (B.1.1.529)-specific antibody shows weak affinity to Delta (B.1.617.2) (Figure 5A)). This challenge that arises when using CFT-SW is most likely to occur when analyzing samples with high sequence homology. Subjecting the Delta strain spike protein mRNA and the Omicron spike protein mRNA to CFT independently and then analyzing by SW shows that an antibody that should be specific to the Omicron (B.1.1.529) spike protein results in weak signal from the Delta (B.1.617.2) sample (Figure 5A) due to antibody cross-reactivity. Similarly, CFT-SW of these same samples using an antibody that should be specific to the Delta (B.1.617.2) spike protein results in signal from the Omicron (B.1.1.529) sample in addition to non-specific binding observed in the blank control sample (Figure 5B). While the mRNA constructs that encode for each of these proteins may be analyzed individually to avoid these challenges, there are instances where analysis of the mRNA mixture may be necessary to better understand the functionality of the mRNA constructs in the presence of one another, such as with co-formulated drug product characterization.

**Figure 5.**
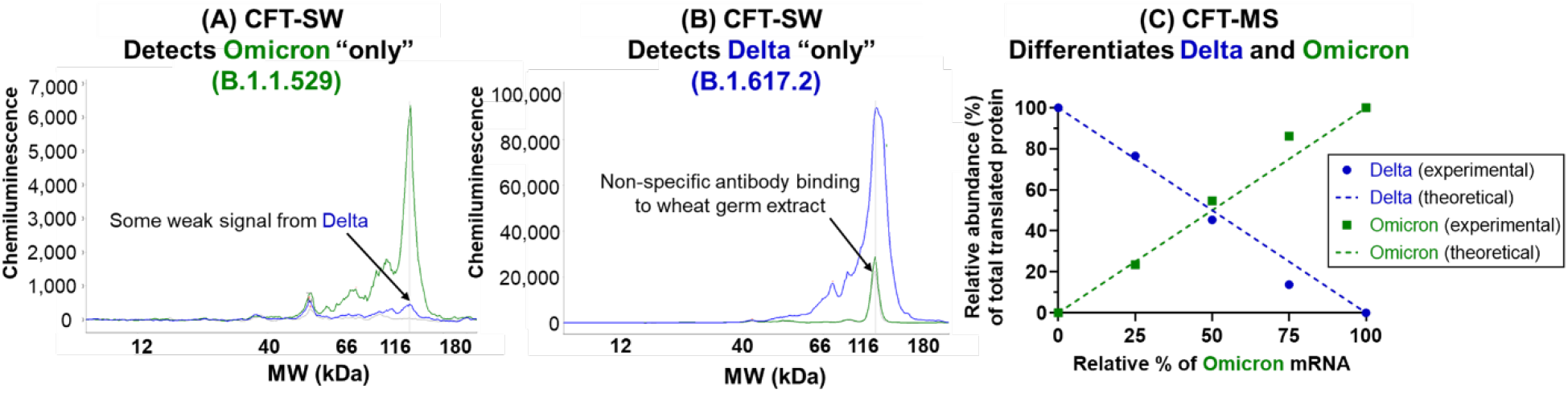
Resulting profiles from CFT-SW of a mixture of SARS-CoV-2 spike protein Delta (B.1.617.2, blue) and Omicron (B.1.1.529, green) strain mRNA using an antibody that should be specific to Omicron (B.1.1.529) only (A) or specific to Delta (B.1.617.2) only (B), demonstrating the cross-reactivity of antibodies that can occur when proteins have high sequence homology (in this case, >97% sequence similarity). Alternatively, analyzing multiple mixtures of Delta (B.1.617.2) and Omicron (B.1.1.529) spike protein mRNA with CFT-MS demonstrates the utility in using mass spectrometry for such a scenario (C). Relative abundance (%) of detected Delta (B.1.617.2) and Omicron (B.1.1.529) spike protein following CFT of mRNA mixtures with varying ratios (solid data points) agrees well with the theoretical values based on mixing ratios (dashed lines).

To combat this challenge, the CFT-MS workflow may be employed. Because MS is a highly selective technique capable of identifying portions of the protein sequence that have unique amino acid combinations even in cases of high overall sequence homology, this antibody-free approach overcomes the multiplexing limitations of cross-reactivity in antibody-based analysis of mRNA mixtures. Mixing the Delta (B.1.617.2) strain spike protein mRNA and the Omicron (B.1.1.529) strain spike protein mRNA at varying ratios (experimental design illustrated in Supplementary Information Figure S5) and then subjecting these mRNA mixtures to CFT-MS, we observe that the amount of translated spike proteins match the theoretical mixing ratios of the mRNA constructs (Figure 5C). This highlights the ability of the platform-based MS detection approach to multiplex protein detection even in cases of high sequence homology, which is particularly advantageous in the analysis of multi-valent mRNA vaccines. It is important to note, however, that different mRNA constructs may have varying translation efficiencies^31,32^ which can result in translated protein relative abundances that do not necessarily reflect the ratio at which the mRNAs were mixed. Because the Delta (B.1.617.2) mRNA and Omicron (B.1.1.529) mRNA used in this work appear to have comparable translation efficiencies, they serve as a clear proof-of-concept for the value of using a platform-based mass spectrometry detection approach to generate relative abundance information of proteins translated from mRNAs, even in mixtures.

### Assessment of mRNA functionality of formulated multi-valent mRNA drug product in human cells by CBT-MS

Using a CFT-MS approach to characterize proteins translated from individual or multi-valent mRNA mixtures following cell-free translation allows for rapid feedback about mRNA quality and functionality in early stages of development without the potentially confounding impact of lipid nanoparticle (LNP) delivery systems as mentioned previously. However, there remain advantages to using a cell-based translation assay such as the ability to interrogate the impact of selected LNP formulations on mRNA translation. Because of this, we tested the feasibility of characterizing proteins generated using a human cell line (Huh7). In this proof-of-concept study, the cells were transfected with two different doses (either 6.67 ng per mRNA or 4.44 ng per mRNA) of a hexavalent mRNA drug product (DP). Each of the six mRNAs in the DP used here encoded for a different spike protein, each with varying degrees of sequence homology (27 to 97%). Following incubation, the cells were lysed and the proteins analyzed using the platform-based MS detection workflow. Results show the confirmation of protein translation from each of the six mRNA constructs (Figure 6).

**Figure 6.**
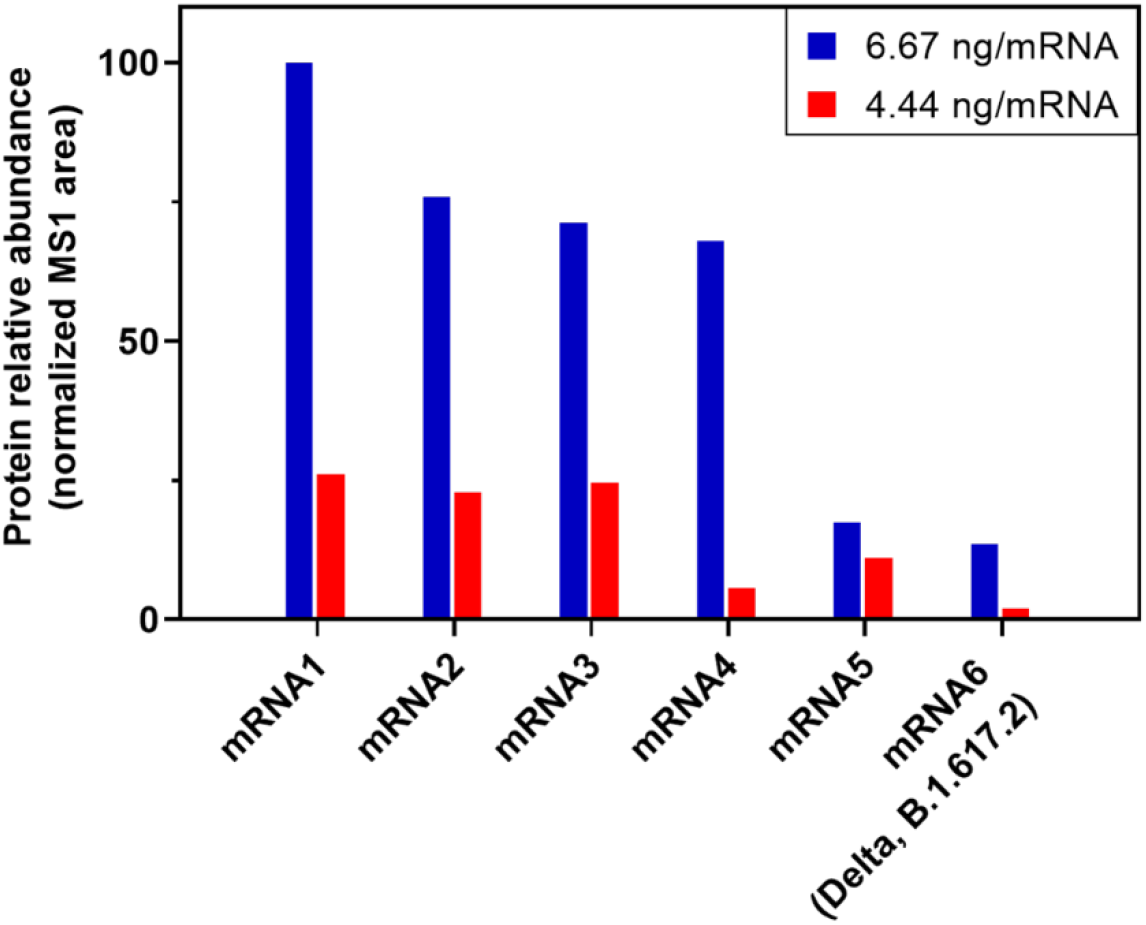
Resulting protein translation from hexavalent mRNA vaccine incubated with Huh7 cells at a dose of 6.67 ng per mRNA construct (blue) or 4.44 ng per mRNA construct. Translated proteins were detected with the LC-MS/MS platform workflow as described here and protein translation is shown as relative abundance (MS1 area). Results show that the proteins are translated in an mRNA construct and dose-dependent manner.

Comparing the relative abundance of each translated protein (represented by MS1 peak area), results show that the cells transfected with 6.67 ng per mRNA produce a greater amount of each protein than those transfected with 4.44 ng per mRNA. This indicates a logical dependence of protein translation on the mRNA dose given to cells and agrees with the CFT-MS dose dependent experiments outlined earlier in this manuscript. Within a given mRNA dosage, e.g., comparing the six translated proteins within the 6.67 ng per mRNA dose, the relative amount of translated protein appear to not be the same. Absolute quantification will be required to confirm the expression differences from multi-valent mRNA to compare mRNA translation efficiency between the constructs. Future work will interrogate the impact of multi-valency on mRNA translation efficiency compared with single valency in addition to establishing cell-based potency assays with high quality antibodies.^33,34^ Ultimately, looking at the neutralized antibodies in mice or non-human primates would provide critical information regarding the impact of multi-valency *in vivo*.

### Assessment of translation fidelity by identifying off-target protein translation and characterizing sites of +1 ribosomal frameshifting with CFT-MS

The use of modified ribonucleotides in therapeutic mRNAs is critical in decreasing innate immunogenicity and improving mRNA stability.^8–10^ Yet despite the advantages, there is still much to be understood about the impact on mRNA translation fidelity. For example, a recent report demonstrated the observation of +1 ribosomal frameshifting at “slippery sites” in N^1^-methylpsuedouridine modified mRNAs. This unintended frameshifting can ultimately change the protein sequence that is translated by the ribosome following the frameshifting junction. Of note, the BNT162b2 SARS-CoV-2 mRNA vaccine that uses N^1^-methylpseudouridine was also shown to induce off-target immunity in both mice and humans to the “mistranslated” protein following vaccination. Despite no evidence to-date of off-target effects from production of the frameshifted protein following vaccination, this work represents a clear example to why it is important to enable analytical detection and characterization of these frameshifted products in an effort to identify such sites that may require further mRNA sequence optimization.

In the Delta (B.1.617.2) spike mRNA used throughout this manuscript, the codon sequence was optimized using an algorithm that did not consider removal of these slippery sequences. Therefore, there are sites that may result in unintended +1 ribosomal frameshifting (Figure 7A). To test whether these sites could be detected from workflows established here, the Delta (B.1.617.2) spike mRNA was subjected to CFT with wheat germ extract and the resulting sample analyzed by the platform-based, untargeted MS detection approach. Analyzing the resulting data by considering both the full in-frame translated protein and one of the possible +1 frameshifted (+1FS) proteins showed a positive identification of both the in-frame and +1FS species. In this instance, the +1FS product was observed at just under 7% relative abundance when comparing the MS1 peak area of the +1FS protein with the in-frame protein, which agrees well with the reported N^1^-methylpseudouridylated firefly luciferase +1FS observed at 8% of the corresponding in-frame protein.^13^

**Figure 7.**
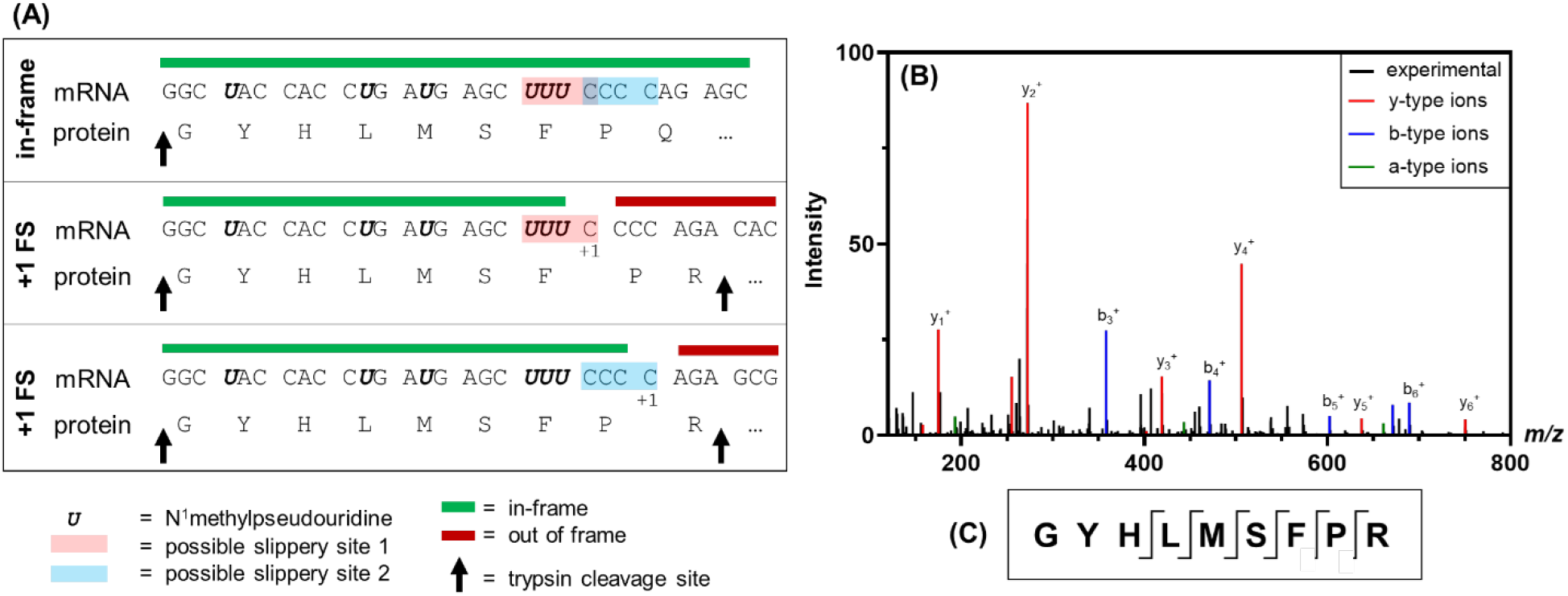
Segment of SARS-CoV-2 Delta (B.1.617.2) spike protein mRNA sequence and corresponding translated protein amino acid sequence for in-frame and +1 ribosomal frameshifted products (A). CFT-MS analysis confirms the presence of the GYHLMSFPR peptide that proves translation of the frameshifted protein (B) *via* comparing the experimentally-observed MS/MS spectrum (black) with theoretical y-(red), b-(blue), and a-type (green) fragments. Additional confirmed fragments such as internal fragments are not labeled. The location of all identified peptide fragments from MS/MS is shown in (C), demonstrating through coverage of the C-terminus where the +1 frameshifted amino acid sequence begins. “+1FS” = +1 ribosomal frameshift.

Because of the low relative abundance of the +1FS product, the sample was again analyzed by CFT-MS but rather than using the platform-based MS method, the peptide GYHLMSFPR, which is indicative of the +1FS product and covers the frameshifting junction, was targeted for fragmentation with no retention time constraints. The resulting data showed high quality MS fragmentation spectra (Figure 7B) with clear confirmation of this peptide at the same retention time it was observed in the untargeted MS analysis. The fragments observed from targeted MS analysis of this peptide (Figure 7C) unequivocally confirm the presence of the +1FS product and, in this case, the amino acid junction at which frameshifting occurs. In this example, a +1FS at the UUU-C motif or the CCC-C motif could result in the same +1FS amino acid sequence upon frameshifted translation. It is possible that one or both of these slippery sites are the root cause of observing the +1FS product, but point mutations of the mRNA sequence would be needed to confirm which is beyond the scope of this work. Of note, the previous report used LC-MS/MS to help narrow in on possible junction sites of +1FS required immunoprecipitation of the translated protein(s) from cell-based translation followed by SDS-PAGE separation and in-gel digestion of the major +1FS species prior to MS analysis, all of which require lengthy sample preparation. Furthermore, that reported workflow in which the major +1FS species were isolated does not consider all possible sites of frameshifting. Advantageously, the CFT-MS workflow demonstrated here enables a more efficient detection approach for assessing +1FS product(s) and can help to identify the corresponding mRNA junction(s).

Given that one of the mRNA constructs used in the CBT-MS experiments (“mRNA 6” in Figure 6) was the same N^1^-methylpseudouridylated Delta (B.1.617.2) spike mRNA analyzed by CFT-MS in Figure 7, the +1 ribosomal frameshifted off-target product was considered when searching the CBT-MS data as well, but was not identified. This is likely due to the significantly lower abundance of the frameshifted product and could not be captured by the untargeted CBT-MS workflow using these mRNA doses as previously reported. Despite this, the CFT-MS approach described above can still provide +1 ribosomal frameshifting feedback and the ability of MS-based detection to provide relative abundance information about the translated proteins from multi-valent mRNA mixtures using either CFT or CBT enables feedback that can advance mRNA vaccines in early stages of development.

## Conclusions

Assessment of protein translation from mRNA provides valuable insights into mRNA functionality and quality. Cell-free translation (CFT) offers rapid sample preparation and a simplified model for protein translation, free from the confounding effects of lipid nanoparticles (LNPs). On the other hand, cell-based translation (CBT) assesses mRNA formulated in LNPs and provides a more representative understanding of mRNA functionality *in vivo*. While each translation method has its unique applications, detecting translated proteins using antibody-based methods can pose challenges due to the expensive and time-consuming antibody screening process as well as the lack of antibody specificity. To address these limitations, a novel platform-based, antibody-free detection method using LC-MS/MS was developed. This approach enables fast and flexible detection of proteins translated from mRNA using either CFT or CBT inputs, making it particularly suitable for rapidly evolving viruses.

To evaluate the effectiveness of the CFT-MS workflow, we employed mRNA encoding for the SARS-CoV-2 Delta (B.1.617.2) spike protein. The results demonstrated that MS not only identified the translated protein, but also provided detailed insights into the primary sequence and relative abundance of the translated protein in an mRNA dose-dependent manner. Additionally, the CFT-SW and CFT-MS workflows were able to assess mRNA quality and integrity following thermal degradation. Further exploration of CFT-MS in the analysis of a bivalent mixture of mRNAs encoding for SARS-CoV-2 Delta (B.1.617.2) spike protein and SARS-CoV-2 Omicron (B.1.1.529) spike protein illustrated the ability of mass spectrometry to multiplex protein detection and highlights the specificity of this technique, even with significant protein sequence homology. We also established a CBT-MS workflow, with the MS component continuing to be platform-based, that demonstrated a dose-dependent response of translated proteins originating from a hexavalent mRNA vaccine drug product.

Furthermore, the utility of CFT-MS is exemplified in the ability to detect +1 ribosomal frameshifted Delta (B.1.617.2) spike protein caused by N^1^-methylpseudouridylation of the mRNA. This highlights the capability of this approach to assess mRNA translation fidelity and provides a framework for identifying regions of mRNA sequences that may require optimization to prevent unintended frameshifting at “slippery sequences.” One of the biggest advantages of mRNA vaccine technology is the rapid development that can enable critical response to quickly evolving diseases, yet this attribute requires that analytical methodologies can keep up. For the first time in peer-reviewed literature, a platform-based, antibody-free mass spectrometry detection method that can be coupled to either cell-free or cell-based mRNA translation was established to provide key information about single- or multi-valent mRNA vaccines such as translated protein identity, mRNA quality, and mRNA translation efficiency and fidelity. These CFT-MS and CBT-MS techniques enable pharmaceutical development to quickly pivot in response to rapidly evolving viruses.

## Author contributions

AQS, BWR, and XL were responsible for overall experimental design. AQS performed sample preparation, data analysis, and interpretation of mass spectrometry experiments; BWR performed sample preparation, data analysis, and interpretation of cell-free translation and Simple Western™ experiments. AQS, BWR, XL, MH, JK, MS, RR, and HS were responsible for cell-free translation experimental design. CT, AG, AQS, and XL were responsible for cell-based translation experimental design. AG performed cell-based translation sample preparation. AQS and BWR were major contributors in writing the manuscript. All authors read and approved the final manuscript.

## Acknowledgments

The authors would like to thank Natalia Marusa, formerly of Eurofins Lancaster Laboratories Professional Service, Lancaster, PA, and currently of Merck & Co., Inc. Rahway, NJ, USA for her contributions to the capillary electrophoresis analysis of mRNA. The authors also appreciate Sonak Dalal of Eurofins Lancaster Laboratories Professional Service, Lancaster, PA, for his assistance with mass spectrometry sample preparation. The authors appreciate helpful scientific discussions with Chris Wang, Dr. Dustin Klein, and Dr. William Cantara regarding the +1 ribosomal frameshifting experiments and with Dr. Josef Vlasak regarding cell-free translation.

## Data availability

The datasets generated and/or analyzed during the current study are not publicly available due to being company confidential information but are available from the corresponding authors on reasonable request.

## References

1. Deviatkin, A. A. et al. Universal Flu mRNA Vaccine: Promises, Prospects, and Problems. Vaccines 10, 709 (2022).

2. Mu, Z., Haynes, B. F. & Cain, D. W. HIV mRNA Vaccines—Progress and Future Paths. Vaccines 9, 134 (2021).

3. Falsey, A. R. et al. Efficacy and Safety of an Ad26.RSV.preF-RSV preF Protein Vaccine in Older Adults. N. Engl. J. Med. 388, 609–620 (2023).

4. Pardi, N., Hogan, M. J. & Weissman, D. Recent advances in mRNA vaccine technology. Curr. Opin. Immunol. 65, 14–20 (2020).

5. Chaudhary, N., Weissman, D. & Whitehead, K. A. mRNA vaccines for infectious diseases: principles, delivery and clinical translation. Nat. Rev. Drug Discov. 20, 817–838 (2021).

6. Pardi, N., Hogan, M. J., Porter, F. W. & Weissman, D. mRNA vaccines — a new era in vaccinology. Nat. Rev. Drug Discov. 17, 261–279 (2018).

7. The Nobel Prize in Physiology or Medicine 2023. NobelPrize.org https://www.nobelprize.org/prizes/medicine/2023/summary/.

8. Anderson, B. R. et al. Incorporation of pseudouridine into mRNA enhances translation by diminishing PKR activation. Nucleic Acids Res. 38, 5884–5892 (2010).

9. Holtkamp, S. et al. Modification of antigen-encoding RNA increases stability, translational efficacy, and T-cell stimulatory capacity of dendritic cells. Blood 108, 4009–4017 (2006).

10. Andries, O. et al. N1-methylpseudouridine-incorporated mRNA outperforms pseudouridine-incorporated mRNA by providing enhanced protein expression and reduced immunogenicity in mammalian cell lines and mice. J. Controlled Release 217, 337–344 (2015).

11. Nance, K. D. & Meier, J. L. Modifications in an Emergency: The Role of N1-Methylpseudouridine in COVID-19 Vaccines. ACS Cent. Sci. 7, 748–756 (2021).

12. Vogel, A. B. et al. BNT162b vaccines protect rhesus macaques from SARS-CoV-2. Nature 592, 283–289 (2021).

13. Mulroney, T. E. et al. N1-methylpseudouridylation of mRNA causes +1 ribosomal frameshifting. Nature 625, 189–194 (2024).

14. Analytical Procedures for mRNA Vaccine Quality (Draft Guidelines)- 2nd Edition | USP-NF. https://www.uspnf.com/notices/analytical-procedures-mrna-vaccines-20230428.

15. Sanyal, G., Särnefält, A. & Kumar, A. Considerations for bioanalytical characterization and batch release of COVID-19 vaccines. NPJ Vaccines 6, 53 (2021).

16. Poveda, C., Biter, A. B., Bottazzi, M. E. & Strych, U. Establishing Preferred Product Characterization for the Evaluation of RNA Vaccine Antigens. Vaccines 7, 131 (2019).

17. Gregorio, N. E., Levine, M. Z. & Oza, J. P. A User’s Guide to Cell-Free Protein Synthesis. Methods Protoc. 2, 24 (2019).

18. Rosenblum, G. & Cooperman, B. S. Engine out of the chassis: cell-free protein synthesis and its uses. FEBS Lett. 588, 261–268 (2014).

19. Zemella, A., Thoring, L., Hoffmeister, C. & Kubick, S. Cell-Free Protein Synthesis: Pros and Cons of Prokaryotic and Eukaryotic Systems. Chembiochem 16, 2420–2431 (2015).

20. Perez, J. G., Stark, J. C. & Jewett, M. C. Cell-Free Synthetic Biology: Engineering Beyond the Cell. Cold Spring Harb. Perspect. Biol. 8, 1–26 (2016).

21. Fogeron, M.-L., Lecoq, L., Cole, L., Harbers, M. & Böckmann, A. Easy Synthesis of Complex Biomolecular Assemblies: Wheat Germ Cell-Free Protein Expression in Structural Biology. Front. Mol. Biosci. 8, (2021).

22. Harbers, M. Wheat germ systems for cell-free protein expression. FEBS Lett. 588, 2762–2773 (2014).

23. Raffaele, J., Loughney, J. W. & Rustandi, R. R. Development of a microchip capillary electrophoresis method for determination of the purity and integrity of mRNA in lipid nanoparticle vaccines. ELECTROPHORESIS 43, 1101–1106 (2022).

24. Hartsough, E. M., Shah, P., Larsen, A. C. & Chaput, J. C. Comparative analysis of eukaryotic cell-free expression systems. BioTechniques 59, 149–151 (2015).

25. Nguyen, U., Squaglia, N., Boge, A. & Fung, P. A. The Simple Western™: a gel-free, blot-free, hands-free Western blotting reinvention. Nat. Methods 8, v–vi (2011).

26. Scheller, C., Krebs, F., Wiesner, R., Wätzig, H. & Oltmann-Norden, I. A comparative study of CE-SDS, SDS-PAGE, and Simple Western—Precision, repeatability, and apparent molecular mass shifts by glycosylation. ELECTROPHORESIS 42, 1521–1531 (2021).

27. Gillespie, P. F. et al. Understanding the Spike Protein in COVID-19 Vaccine in Recombinant Vesicular Stomatitis Virus (rVSV) Using Automated Capillary Western Blots. ACS Omega 8, 3319–3328 (2023).

28. Sutton, W. J. H. et al. Quantification of SARS-CoV-2 spike protein expression from mRNA vaccines using isotope dilution mass spectrometry. Vaccine 41, 3872–3884 (2023).

29. Ozdilek, A., Paschall, A. V., Dookwah, M., Tiemeyer, M. & Avci, F. Y. Host protein glycosylation in nucleic acid vaccines as a potential hurdle in vaccine design for nonviral pathogens. Proc. Natl. Acad. Sci. 117, 1280–1282 (2020).

30. Chalkias, S. et al. Safety, immunogenicity and antibody persistence of a bivalent Beta-containing booster vaccine against COVID-19: a phase 2/3 trial. Nat. Med. 28, 2388–2397 (2022).

31. Boël, G. et al. Codon influence on protein expression in E. coli correlates with mRNA levels. Nature 529, 358–363 (2016).

32. Kim, S. C. et al. Modifications of mRNA vaccine structural elements for improving mRNA stability and translation efficiency. Mol. Cell. Toxicol. 18, 1–8 (2022).

33. Li, H. H. et al. Development and qualification of cell-based relative potency assay for a human respiratory syncytial virus (RSV) mRNA vaccine. J. Pharm. Biomed. Anal. 234, 115523 (2023).

34. Patel, N. et al. Development and Characterization of an In Vitro Cell-Based Assay to Predict Potency of mRNA–LNP-Based Vaccines. Vaccines 11, 1224 (2023).

