## Supplementary Information for "Enabling functionality and translation fidelity characterization of mRNA-based vaccines with a platform-based, antibody-free mass spectrometry detection approach"

|  | Total | Mass only | Mass and MS/MS |
|---|---|---|---|
| Delta spike_combined coverage | 99.1% | 5.2% | 93.9% |
| Delta spike_trypsin | 67.2% | 10.9% | 56.3% |
| Delta spike_alpha lytic protease | 90.1% | 34.0% | 56.1% |
| Delta spike_chymotrypsin | 98.7% | 17.5% | 81.2% |

```
........10 ........20 ........30 ........40 ........50 ........60 ........70 ........80
MFVFLVLLPL VSSQCVNLRT RTQLPPAYTN SFTRGVYYPD KVFRSSVLHS TQDLFLPFFS NVTWFHAIHV SGTNGTKRFD Combined coverage
MFVFLVLLPL VSSQCVNLRT RTQLPPAYTN SFTRGVYYPD KVFR                                   RFD Trypsin
MFVFLVLLPL VS QCVNLRT RTQLPPAYTN SFTRGVYYPD KVFRS VLHS TQDLFLPFFS NVTWFHAIHV S  NGTKRFD Alpha lytic protease
MFVFLVLLPL VSSQCVNLRT RTQLPPAYTN SFTRGVYYPD KVFRSSVLHS TQDLFLPFFS NVTWFHAIHV SGTNGTKRFD Chymotrypsin

.......90 .......100 .......110 .......120 .......130 .......140 .......150 .......160
NPVLPFNDGV YFASTEKSNI IRGWIFGTTL DSKTQSLLIV NNATNVVIKV CEFQFCNDPF LDVYYHKNNK SWMESGVYSS Combined coverage
NPVLPFNDGV YFASTEKSNI IRGWIFGTTL DSKTQSLLIV NNATNVVIKV CEFQFCNDPF LDVYYHKNNK SWMESGVYSS Trypsin
NPVLPFNDGV YFAS EKSNI IRGWIFGTTL DSKTQSLLIV NNA NVV    CEFQFCNDPF LDVYYHKNNK SWMESGVYS  Alpha lytic protease
NPVLPFNDGV YFASTEKSNI IRGWIFGTTL DSKTQSLLIV NNATNVVIKV CEFQFCNDPF L YHKNNK   SWMESGVYSS Chymotrypsin

.......170 .......180 .......190 .......200 .......210 .......220 .......230 .......240
ANNCTFEYVS QPFLMDLEGK QGNFKNLREF VFKNIDGYFK IYSKHTPINL VRDLPQGFSV LEPLVDLPIG INITRFQTLL Combined coverage
ANNCTFEYVS QPFLMDLEGK QGNFKNLREF VFKNIDGYFK IYSKHTPINL VRDLPQGFSV LEPLVDLPIG INITRFQTLL Trypsin
 NNCTFEYVS QPFLMDLEGK QGNFKNLREF VFKNIDGYFK IYSKHTPINL VRDLPQGFSV LEPLVDLPIG INITRFQTLL Alpha lytic protease
ANNCTFEYVS QPFLMDLEGK QGNFKNLREF VFKNIDGYFK IYSKHTPINL VRDLPQGFSV LEPLVDLPIG INITRFQTLL Chymotrypsin

.......250 .......260 .......270 .......280 .......290 .......300 .......310 .......320
ALHRSYLTPG DSSSGWTAGA AAYYVGYLQP RTFLLKYNEN GTITDAVDCA LDPLSETKCT LKSFTVEKGI YQTSNFRVQP Combined coverage
ALHRSYLTPG DSSSGWTAGA AAYYVGYLQP RTFLLKYNEN GTITDAVDCA LDPLSETKCT LKSFTVEKGI YQTSNFRVQP Trypsin
ALHRSYLTPG DS  GWTA    AYYVGYLQP RTFLLKYNEN GTIT   DCA LDPLSETKCT LKSFTVEKGI YQTSNFRVQP Alpha lytic protease
ALHRSYLTPG DSSSGWTAGA AAYYVGYLQP RTFLLKYNEN GTITDAVDCA LDPLSETKCT LKSFTVEKGI YQTSNFRVQP Chymotrypsin

.......330 .......340 .......350 .......360 .......370 .......380 .......390 .......400
TESIVRFPNI TNLCPFGEVF NATRFASVYA WNRKRISNCV ADYSVLYNSA SFSTFKCYGV SPTKLNDLCF TNVYADSFVI Combined coverage
TESIVRFPNI TNLCPFGEVF NATRFASVYA WNRKRISNCV ADYSVLYNSA SFSTFKCYGV SPTKLNDLCF TNVYADSFVI Trypsin
TESIVRFPNI TNLCPFGEVF NATRFA  YA WNRKRISNCV     VLYNSA    TFKCYGV SPTKLNDLCF TNV  DSFVI Alpha lytic protease
TESIVRFPNI TNLCPFGEVF NATRFASVYA WNRKRISNCV ADYSVLYNSA SFSTFKCYGV SPTKLNDLCF TNVYADSFVI Chymotrypsin

.......410 .......420 .......430 .......440 .......450 .......460 .......470 .......480
RGDEVRQIAP GQTGKIADYN YKLPDDFTGC VIAWNSNNLD SKVGGNYNYR YRLFRKSNLK PFERDISTEI YQAGSKPCNG Combined coverage
RGDEVRQIAP GQTGKIADYN YKLPDDFTGC VIAWNSNNLD SKVGGNYNYR YRLFRKSNLK PFER                  Trypsin
RGDEVRQIAP GQTGKIADYN YKLPDDFTGC VIAWNSNNLD SKVGGNYNYR YRLFRKSNLK PFERDISTEI YQAGSKPCNG Alpha lytic protease
RGDEVRQIAP GQTGKIADYN YKLPDDFTGC VIAWNSNNLD SKVGGNYNYR YRLFRKSNLK PFERDISTEI YQAGSKPCNG Chymotrypsin

.......490 .......500 .......510 .......520 .......530 .......540 .......550 .......560
VEGFNCYFPL QSYGFQPTNG VGYQPYRVVV LSFELLHAPA TVCGPKKSTN LVKNKCVNFN FNGLTGTGVL TESNKKFLPF Combined coverage
                             VVV LSFELLHAPA TVCGPKKSTN LVKNKCVNFN FNGLTGTGVL TESNKKFLPF Trypsin
VEGFNCYFPL QSYGFQPTNG VGYQPYRVV  LSFELLHAPA  VCGPKKSTN LVKNKCVNFN FNGLTGTGVL TESNKKFLPF Alpha lytic protease
VEGFNCYFPL QSYGFQPTNG VGYQPYRVVV LSFELLHAPA TVCGPKKSTN LVKNKCVNFN FNGLTGTGVL TESNKKFLPF Chymotrypsin

.......570 .......580 .......590 .......600 .......610 .......620 .......630 .......640
QQFGRDIADT TDAVRDPQTL EILDITPCSF GGVSVITPGT NTSNQVAVLY QGVNCTEVPV AIHADQLTPT WRVYSTGSNV Combined coverage
QQFGRDIADT TDAVR                                                              VYSTGSNV Trypsin
QQFGRDIADT    VRDPQTL EILDITPCSF GGV  ITPGT NT NQV VLY QGVNCTEVPV AIHADQLTPT WRVYS GSNV Alpha lytic protease
QQFGRDIADT TDAVRDPQTL EILDITPCSF GGVSVITPGT NTSNQVAVLY QGVNCTEVPV AIHADQLTPT WRVYSTGSNV Chymotrypsin

.......650 .......660 .......670 .......680 .......690 .......700 .......710 .......720
FQTRAGCLIG AEHVNNSYEC DIPIGAGICA SYQTQTNSRR RARSVASQSI IAYTMSLGAE NSVAYSNNSI AIPTNFTISV Combined coverage
FQTR                                                                                   Trypsin
FQTRAGCLIG AEHVNNSYEC DIPIGAGICA SYQTQTNSRR RARS   QSI IAYTMSLGAE NS    NNSI AIPTNFTIS  Alpha lytic protease
FQTRAGCLIG AEHVNNSYEC DIPIGAGICA SYQTQTNSRR RARSVASQSI IAYTMSLGAE NSVAYSNNSI AIPTNFTISV Chymotrypsin

.......730 .......740 .......750 .......760 .......770 .......780 .......790 .......800
TTEILPVSMT KTSVDCTMYI CGDSTECSNL LLQYGSFCTQ LNRALTGIAV EQDKNTQEVF AQVKQIYKTP PIKDFGGFNF Combined coverage
                                               ALTGIAV EQDKNTQEVF AQVKQIYKTP PIKDFGGFNF Trypsin
 TEILPVSMT KTS    MYI CGDS ECSNL LLQYGSFCTQ LNRALTGIAV EQDKNTQEVF AQVKQIYKTP PIKDFGGFNF Alpha lytic protease
TTEILPVSMT KTSVDCTMYI CGDSTECSNL LLQYGSFCTQ LNRALTGIAV EQDKNTQEVF AQVKQIYKTP PIKDFGGFNF Chymotrypsin

.......810 .......820 .......830 .......840 .......850 .......860 .......870 .......880
SQILPDPSKP SKRSFIEDLL FNKVTLADAG FIKQYGDCLG DIAARDLICA QKFNGLTVLP PLLTDEMIAQ YTSALLAGTI Combined coverage
SQILPDPSKP SKRSFIEDLL FNKVTLADAG FIKQYGDCLG DIAARDLICA QK                             Trypsin
SQILPDPSKP SKRSFIEDLL FNKVT  DAG FIKQYGDCLG DIAARDLICA QKFNGLTVLP PLLTDEMIAQ YT  LLAGTI Alpha lytic protease
SQILPDPSKP SKRSFIEDLL FNKVTLADAG FIKQYGDCLG DIAARDLICA QKFNGLTVLP PLLTDEMIAQ YTSALLAGTI Chymotrypsin

.......890 .......900 .......910 .......920 .......930 .......940 .......950 .......960
TSGWTFGAGA ALQIPFAMQM AYRFNGIGVT QNVLYENQKL IANQFNSAIG KIQDSLSSTA SALGKLQNVV NQNAQALNTL Combined coverage
                         FNGIGVT QNVLYENQKL IANQFNSAIG KIQDSLSSTA SALGKLQNVV NQNAQALNTL Trypsin
TSGWTFGA   ALQIPFAMQM AYRFNGIGVT QNVLYENQKL IANQFNSAIG KIQDSLS     ALGKLQNVV NQNAQALNTL Alpha lytic protease
TSGWTFGAGA ALQIPFAMQM AYRFNGIGVT QNVLYENQKL IANQFNSAIG KIQDSLSSTA SALGKLQNVV NQNAQALNTL Chymotrypsin
```

```
.......970 .......980 .......990 ......1000 ......1010 ......1020 ......1030 ......1040
VKQLSSNFGA ISSVLNDILS RLDPPEAEVQ IDRLITGRLQ SLQTYVTQQL IRAAEIRASA NLAATKMSEC VLGQSKRVDF Combined coverage
VKQLSSNFGA ISSVLNDILS RLDPPEAEVQ IDRLITGRLQ SLQTYVTQQL IRAAEIRASA NLAATKMSEC VLGQSKRVDF Trypsin
VKQLSSNFGA IS VLNDILS RLDPPEAEVQ IDRLITGRLQ SLQTYVTQQL IRAAEIRAS       KMSEC VLGQSKRVDF Alpha lytic protease
VKQLSSNFGA ISSVLNDILS RLDPPEAEVQ IDRLITGRLQ SLQTYVTQQL IRAAEIRASA NLAATKMSEC VLGQSKRVDF Chymotrypsin

......1050 ......1060 ......1070 ......1080 ......1090 ......1100 ......1110 ......1120
CGKGYHLMSF PQSAPHGVVF LHVTYVPAQE KNFTTAPAIC HDGKAHFPRE GVFVSNGTHW FVTQRNFYEP QIITTDNTFV Combined coverage
CGK                               NFTTAPAIC HDGKAHFPRE GVFVSNGTHW FVTQR                 Trypsin
CGKGYHLMSF PQSAPHGVVF LHVTYVPAQE KNFTT PAIC HDGKAHFPRE GVFVSNGTHW FVTQRNFYEP QIITTDNTFV Alpha lytic protease
CGKGYHLMSF PQSAPHGVVF LHVTYVPAQE KNFTTAPAIC HDGKAHFPRE GVFVSNGTHW FVTQRNFYEP QIITTDNTFV Chymotrypsin

......1130 ......1140 ......1150 ......1160 ......1170 ......1180 ......1190 ......1200
SGNCDVVIGI VNNTVYDPLQ PELDSFKEEL DKYFKNHTSP DVDLGDISGI NASVVNIQKE IDRLNEVAKN LNESLIDLQE Combined coverage
                             EEL DKYFKNHTSP DVDLGDISGI NASVVNIQKE IDRLNEVAKN LNESLIDLQE Trypsin
S     VIGI VNNTVYDPLQ PELDSFKEEL DKYFKNHTSP DVDLGDISGI NAS VNIQKE IDRLNEVAKN LNES       Alpha lytic protease
SGNCDVVIGI VNNTVYDPLQ PELDSFKEEL DKYFKNHTSP DVDLGDISGI NASVVNIQKE IDRLNEVAKN LNESLIDLQE Chymotrypsin

......1210 ......1220 ......1230 ......1240 ......1250 ......1260 ......1270 1271
LGKYEQYIKW PWYIWLGFIA GLIAIVMVTI MLCCMTSCCS CLKGCCSCGS CCKFDEDDSE PVLKGVKLHY T Combined coverage
LGKYEQYIK                                                 FDEDDSE PVLKGVKLHY T Trypsin
                      GLIAIVMVTI MLCCMTS                            E PVLKGVKLHY T Alpha lytic protease
LGKYEQYIKW PWYIWLGFIA GLIAIVMVTI MLCCMTSCCS CL                 DEDDSE PVLKGVKLHY   Chymotrypsin
```

**Figure S1.** Sequence coverage of SARS-CoV-2 Delta (B.1.617.2) spike protein based on enzymatic digestion with trypsin, chymotrypsin, or alpha lytic protease. Amino acids confirmed with MS/MS matches are represented in green and amino acids identified with mass only matches are represented in blue. Black amino acids indicate no mass only or MS/MS matches were identified.

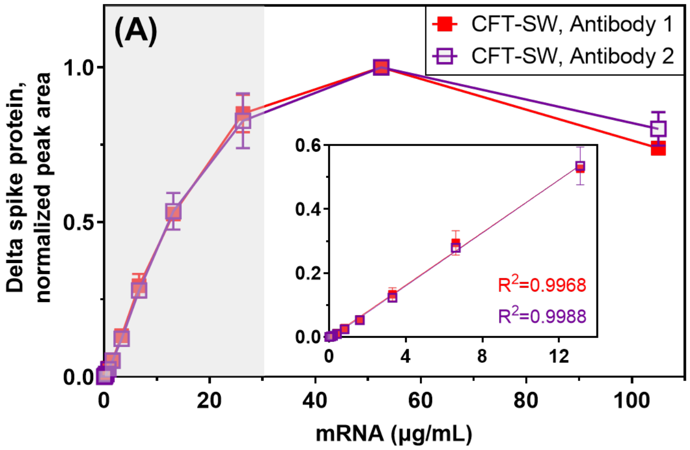

**Figure S2.** Dose response curves of CFT-SW using Antibody 1 (more sensitive, red, Sino Biological) and Antibody 2 (less sensitive, purple, Merck & Co., Inc., Rahway, NJ, USA IP). Y-axis represents normalized peak area within the respective antibody detection.

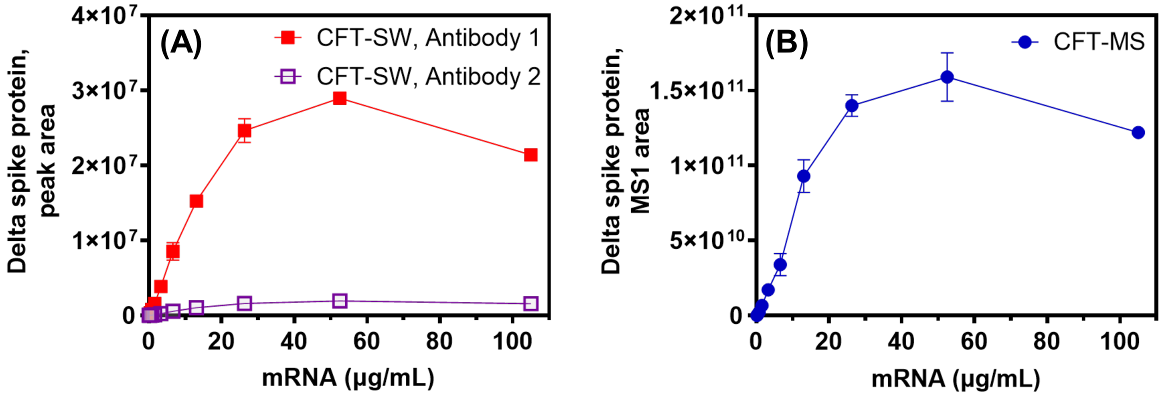

**Figure S3.** Dose response curves of CFT-SW (A) and CFT-MS (B) with the actual (not normalized) peak area (CFT-SW) and MS1 area (CFT-MS). Error bars are shown as %RSD for all data points but at times are obscured by data point size.

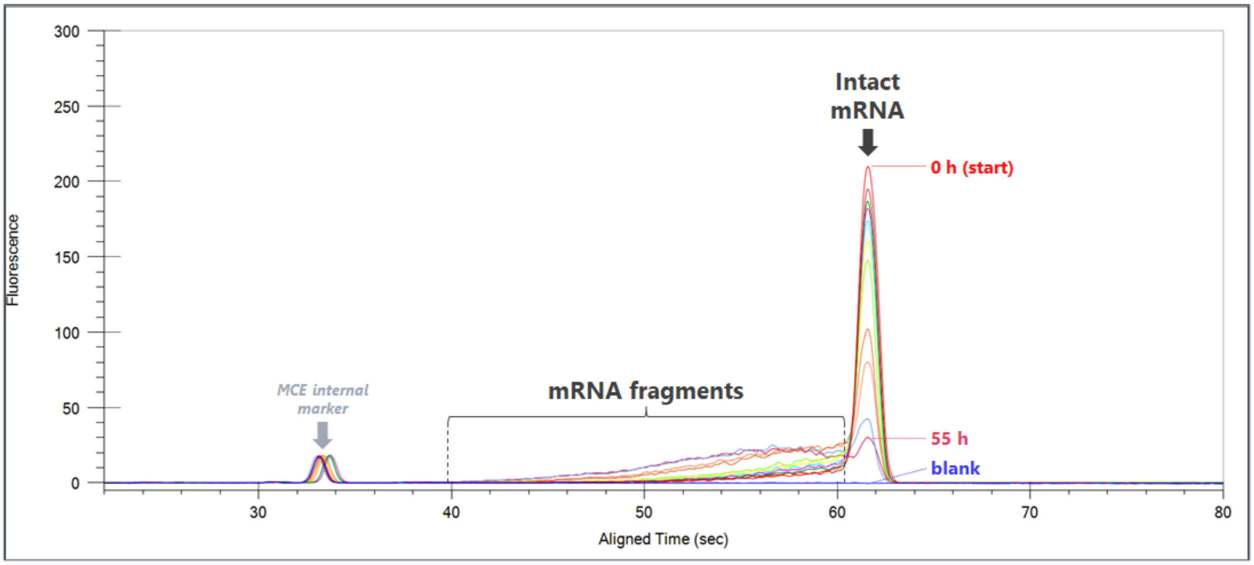

**Figure S4.** Microchip capillary electrophoresis (MCE) electropherograms of SARS-CoV-2 Delta (B.1.617.2) mRNA degraded at 50 °C. MCE diluent without mRNA was used as a blank blue trace). Electropherograms were aligned according to the intact mRNA peak.

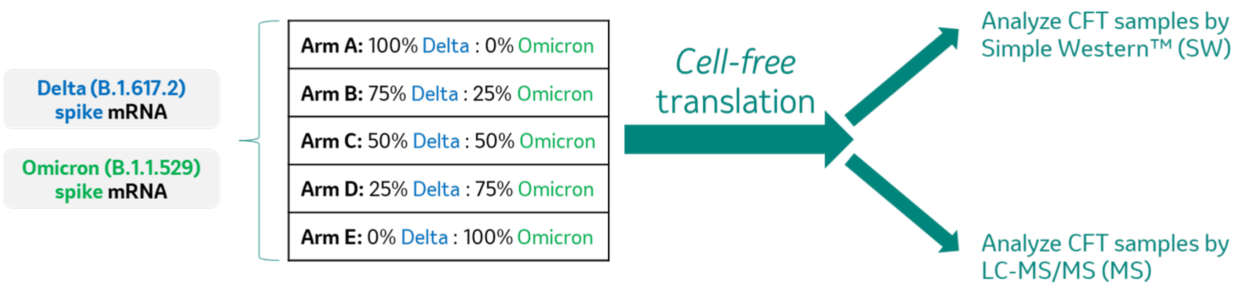

**Figure S5.** Schematic showing the mixing scheme of the Delta (B.1.617.2) strain spike protein mRNA and the Omicron (B.1.1.529) strain spike protein mRNA at varying ratios. Results from CFT-MS analysis are shown in the main text Figure 5C.
